# *MLH1/3* variants causing aneuploidy, pregnancy loss, and premature reproductive aging

**DOI:** 10.1101/2021.01.14.426654

**Authors:** Priti Singh, Robert Fragoza, Cecilia S. Blengini, Tina N. Tran, Gianno Pannafino, Najla Al-Sweel, Kerry J. Schimenti, Karen Schindler, Eric A. Alani, Haiyuan Yu, John C. Schimenti

## Abstract

Most spontaneous pregnancy losses are a result of embryonic aneuploidy stemming from mis-segregation of chromosomes during meiosis. Proper disjunction of homologous chromosomes is dependent upon precise control of crossing-over, a process requiring the mismatch repair (MMR) genes *MLH1* and *MLH3*. Both are required for fertility and completion of meiosis in mice. People inheriting variants in these genes are often at high risk for colorectal cancer and Lynch syndrome, yet the potential impacts of variants upon reproduction are unclear. To determine if *MLH1/3* variants (namely single nucleotide polymorphisms, or SNPs) in human populations can cause reproductive abnormalities, we used a combination of computational predictions, yeast two-hybrid assays, and assays of MMR and recombination in yeast to select nine *MLH1* and *MLH3* variants for modeling in mice via genome editing. We identified 7 alleles that caused reproductive defects in mice including subfertility in females, male infertility, reduced sperm counts, and increased spermatocyte apoptosis. Remarkably, these alleles in females caused age-dependent decreases in litter size, and increased resorption of embryos during pregnancy. These outcomes were likely a consequence of reduced meiotic chiasmata, in turn causing an increase in misaligned chromosomes and univalents in meiotic metaphase I (MI). Our data indicate that segregating hypomorphic alleles of meiotic recombination genes in populations can predispose females to increased incidence of pregnancy loss from gamete aneuploidy.

## Introduction

The essential function of meiotic recombination is to drive pairing of homologous chromosomes, ensuring proper disjunction of homologs to daughter cells at the first meiotic division. Failure of a chromosome pair to undergo at least one crossover per chromosome arm predisposes to formation of aneuploid gametes that, if fertilized, leads to pregnancy loss. Approximately 50-70% of early miscarriages are associated with chromosome abnormalities, with most having aneuploidy, primarily trisomies ^1,2^. It is well recognized that the incidence of aneuploidy in abortuses increases with female age and has been hypothesized to be attributable to non-genetic factors such as decay of chromosome cohesion ^3,4^. However, there are indications that parental genetic factors can predispose to generation of aneuploid conceptuses, and may contribute to cases of recurrent pregnancy loss (RPL; defined as three or more consecutive miscarriages) ^5,6^. For example, much attention has focused on the potential effects of variants/mutations in the synaptonemal complex protein SYCP3 in human RPL ^7,8^, because SYCP3-deficient female mice experience loss of embryos resulting from aneuploid oocytes ^9^. However, it is unclear whether, or to what extent, mutations or deleterious variants in *Sycp3* or other genes cause miscarriages or RPL.

Chromosome mis-segregation during meiosis can occur at Meiosis I (MI) or Meiosis II (MII), though a lack of recombination on extra/missing chromosomes has been associated with inefficient crossover maturation before MI ^10^. The resulting gametes, if fertilized, can lead to aneuploid embryos that are almost universally incompatible with viability, thus causing miscarriage. Several genes are required for crossover (CO) recombination (and thus fertility) in mice, including the mismatch repair (MMR) proteins MLH1 and MLH3, which form a heterodimeric endonuclease required to resolve double Holliday junction recombination intermediates ^11–19^. Null alleles of *Mlh1 or Mlh3* cause meiotic arrest and sterility in mice because they are needed for ~90% of all crossovers (those known as interference-dependent Class I crossovers) ^11,20–25^. The absence of COs leads to univalent chromosomes that fail to align properly while attached to microtubules at the metaphase plate, thus triggering the spindle assembly checkpoint (SAC) and blocking anaphase progression ^26^. However, recombination defects affecting bivalent formation of only one or a few chromosome pairs is inefficient at triggering the SAC, allowing such gametes to survive and thus predisposing to aneuploidy ^27–30^.

We hypothesized that human population variants in *MLH1/3* might cause defects in chromosome segregation during MI or MII, leading to infertility, miscarriage, and/or developmental defects. To test this and gain a comprehensive understanding of the functional impact of such variants, we systematically evaluated human missense single nucleotide polymorphisms (SNPs) curated in the ExAC and (subsequently) gnomAD databases ^31,32^. Minor alleles predicted to be deleterious *in silico* by various algorithms, and which deviated from normal function in biochemical assays or genetic assays in yeast, were selected for modeling in mice. Most of the mutant mouse models were fertile but exhibited crossing over defects that led to age-related decreases in female fecundity, elevated aneuploidy, and consequent increases in pregnancy loss. These results have implications especially for older couples, or those experiencing RPL, who may be at increased risk for having conceptuses or children with trisomies as a result of bearing variants in these recombination genes.

## Results

### Strategy and selection of candidate pathogenic MUTL homolog variants

To test the possibility that segregating infertility variants of *MLH1* or *MLH3* exist in human populations, we identified potentially pathogenic missense SNPs in these genes using methods and criteria outlined in Figure 1A, including: 1) functional prediction algorithms scoring the variant as deleterious; 2) has an allele frequency (AF) within the range of 0.02% to 2% in any gnomAD-listed population; 3) disruptive to binary proteinprotein-interactions in yeast two-hybrid (Y2H) assay; or 4) causes an MMR and/or recombination phenotype in baker’s yeast (*S. cerevisiae*) when the orthologous amino acid is altered. These steps were used to prioritize a subset of *MLH1* and *MLH3* variants for subsequent modeling in mice.

**Figure 1.**
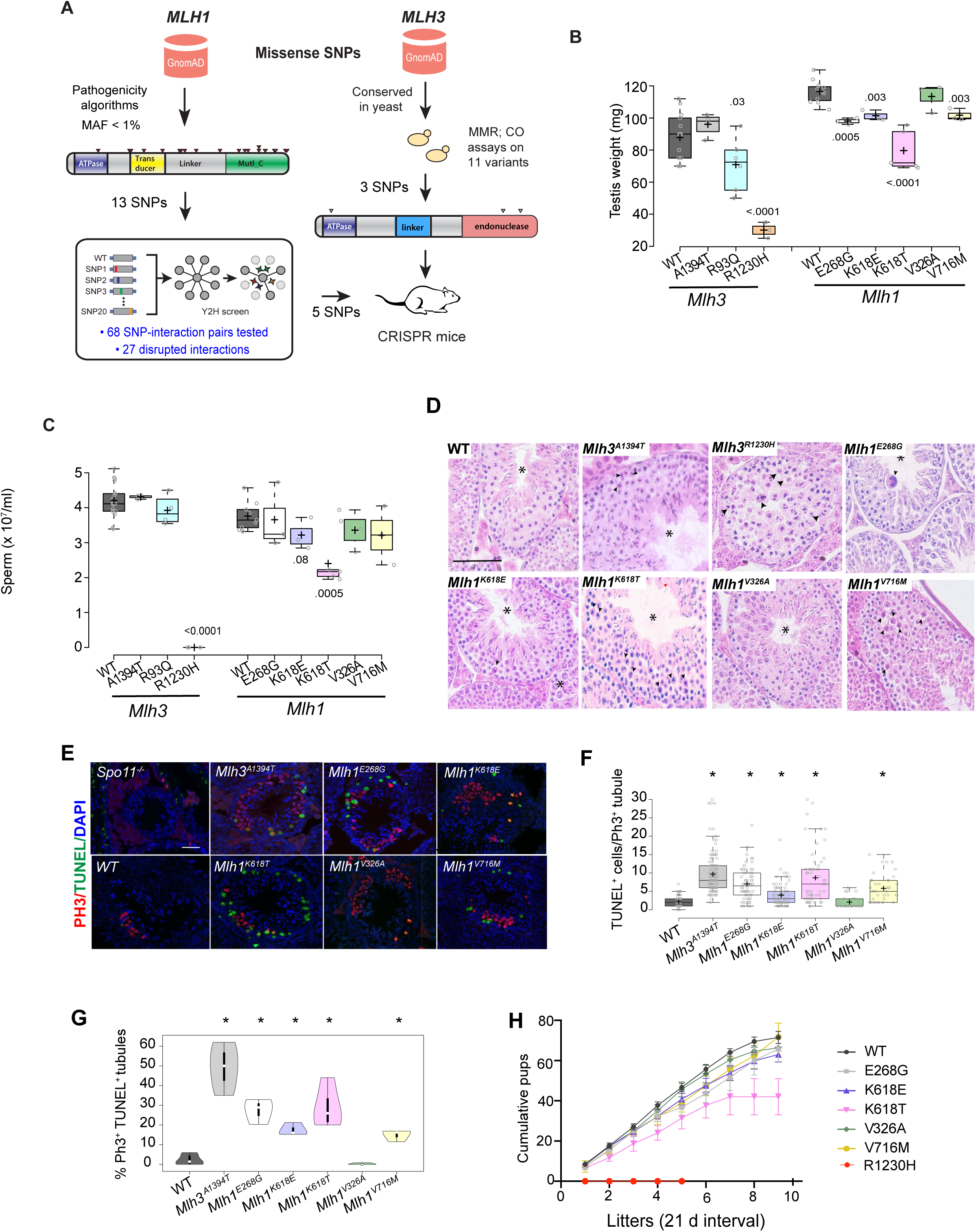
Spermatogenesis defects in mutant mice. **A)** Overview of process for selecting human *MLH1* and *MLH3* variants for mouse modeling. gnomAD (https://gnomad.broadinstitute.org) was the source of allele frequencies. MMR, mismatch repair; CO, crossover. The SNPs tested are listed in Table S1. **B)** Testis weights of mutants. The *Mlh1* and *Mlh3* samples were taken from 6 month and 2-3 month old mice, respectively, hence the difference between WT samples. Each data point is one testis from one animal. Horizontal line in each box is the median, and “+” is the mean. The extreme upper and lower whiskers are the data ranges, and the box lengths are span the 1^st^ and 3^rd^ quartiles. For this and panel C, numbers are p values relative to corresponding WT of that cohort; only genotypes with values <0.05 are shown. Genotype abbreviations: EG, *Mlh1^E268G/E268G^*; KE, *Mlh1^K618E/K618E^*; KT, *Mlh1^K618T/K618T^*; V326A, *Mlh1^V326A/V326A^*; V716M, *Mlh1^V716M/V716M^*. AT, *Mlh3^A1394T/ A1394T^*; RQ, *Mlh3^R93Q/R93Q^*; RH, *Mlh3^R1230H/R1230H^*. **C)** Epididymal sperm counts. All were from 6-month old mice, except *Mlh3^A1394T/ A1394T^* and *Mlh3^R93G/R93G^*, which were from 3 month old animals) Controls were from pooled littermates. Genotype abbreviations are as in “B.” **D)** Testis histology. Examples of H&E stained seminiferous tubule cross sections of homozygotes for the indicated genotypes. Asterisk (*) represents Stage XI-XII tubules with metaphase cells. Arrowheads represent degenerating metaphase cells. Scale Bar = 50 μm. **E)** Testis sections immunolabeled by phosphorylated histone H3 (PH3), and also TUNEL-stained for apoptosis counterstained with DAPI (for DNA). *Mlh1/3* testes are from homozygous mutants. Scale Bar = 50μm. Genotypes are indicated. **F-G)** Quantification of TUNEL data from samples in “E.” Plotted are overall percentages of TUNEL^+^ cells (F) and also tubules containing >4 TUNEL^+^ cells (G). This number was chosen because no WT tubule sections had >4. N≥3 animals. *, p ≤ 0.0003. **H)** Fertility trials of homozygous males and control littermates. *, p ≤ 0.0001. N = 3-5 males per genotype. The X axis values are in 21-day intervals, which approximates litter intervals for fertile females housed continuously with a male. All p values were from Student’s two-tailed t-test. In B,C,F-H, each plot is defined as follows: Horizontal line in box = median; box width = data points between the first and third quartiles of the distribution; whiskers = minimum and maximum values; ‘+’ sign = mean value.

Disease-associated mutations in *MLH1* often function molecularly by disrupting corresponding protein-protein interactions ^33,34^. As such, we searched for all potentially functional *MLH1* missense variants that matched our defined 0.02% < AF < 2% criteria range and then cloned and tested these variants for corresponding disruptions to protein-protein interactions by the Y2H assay. This resulted in 13 *MLH1* variants tested by Y2H across 68 total SNP-interaction pairs. We then further filtered these variants for SNPs that scored as deleterious across multiple functional prediction algorithms. This resulted in five *MLH1* predicted-deleterious variants that disrupted 27 out of 37 (73%) of their corresponding SNP-interactions pairs (Table1; Table S1).

For *MLH3*, we modeled eleven human SNPs in yeast by constructing strains bearing the cognate amino acid (AA) changes in the endogenous *MLH3* gene, then measuring MMR using a *lys2* frameshift reversion assay ^35^. Most of the AAs are located in conserved endonuclease and ATP binding domains (Table S2; Fig. S2). To measure meiotic crossing over, these strains were mated to an *mlh3*Δ strain so that the diploids contained fluorescence markers separated by ~20 cM on chromosome VIII ^36^. The diploids were sporulated to identify, in complete tetrads, presence or absence of crossovers (Fig. S1). Three of the *MLH3* SNPs (rs781779034, rs138006166 and rs781739661) had null-like or intermediate phenotypes for MMR, and null-like crossing over defects (Table 1; Tables S2,S3). Additionally, all three were classified as deleterious by functional prediction algorithms (Table 1). Overall, nine of the eleven alleles modeled in yeast conferred null-like or intermediate phenotypes in either or both tested metrics (Table S2).

**Table 1.**
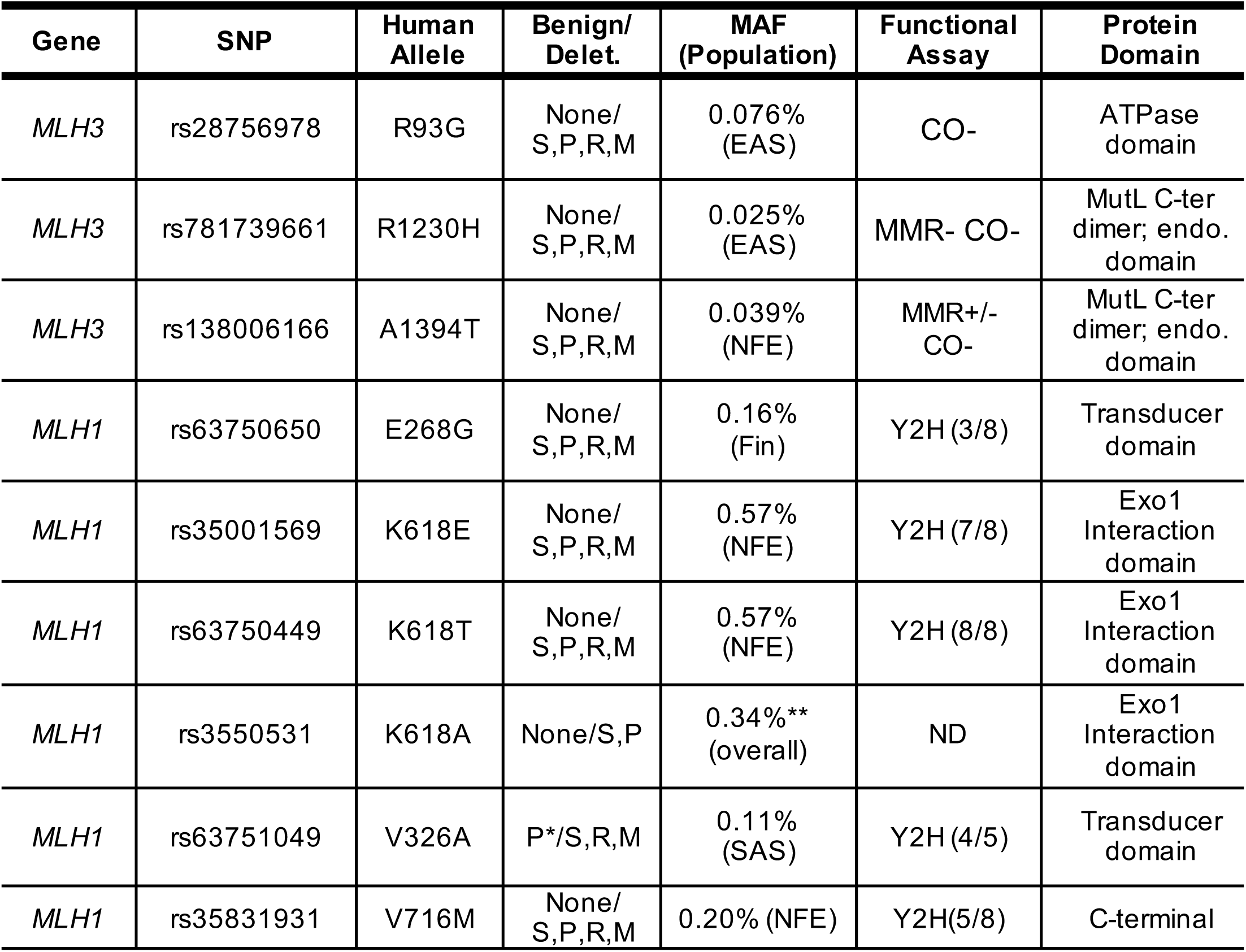
SNPs modeled in mice. gnomAD (http://gnomad.broadinstitute.org) allele frequencies (version 2.1.1) correspond to the population with the highest allele frequency. EAS = East Asian, NFE = European (non-Finnish), Fin = Finnish, SAS = South Asian; S = SIFT; P = Polyphen2; REVEL = R; M = Mutation Assessor. ND = no data; MMR = mismatch repair; CO = meiotic crossing over phenotypes (+, wildtype, -, null); dimer = dimerization; endo = endonuclease; “Y2H” = yeast two-hybrid disruption, with the values in parentheses indicating the fraction of known interactions with proteins disrupted by the corresponding *MLH1* mutation (see details in Table S1. *This pathogenicity score is classified as “possibly” damaging. **This allele was generated near the end of the study, and consists of a dinucleotide change, each corresponding to one of the nucleotides altered in the K618E and K618T alleles. REVEL, Mutation Assessor and CADD scores were not obtained for K618A because this allele is not a recognized population variant. gnomAD lists overall AF for K618A, but its NFE-specific AF is likely identical to K618E and K618T, which almost always co-occur (see text). For functional assays in yeast, see details in Tables S2-S3.

| Gene | SNP | Human Allele | Benign/ Delet. | MAF (Population) | Functional Assay | Protein Domain |
| --- | --- | --- | --- | --- | --- | --- |
| *MLH3* | rs28756978 | R93G | None/ S,P,R,M | 0.076% (EAS) | CO- | ATPase domain |
| *MLH3* | rs781739661 | R1230H | None/ S,P,R,M | 0.025% (EAS) | MMR- CO- | MutL C-ter dimer; endo. domain |
| *MLH3* | rs138006166 | A1394T | None/ S,P,R,M | 0.039% (NFE) | MMR+/- CO- | MutL C-ter dimer; endo. domain |
| *MLH1* | rs63750650 | E268G | None/ S,P,R,M | 0.16% (Fin) | Y2H (3/8) | Transducer domain |
| *MLH1* | rs35001569 | K618E | None/ S,P,R,M | 0.57% (NFE) | Y2H (7/8) | Exo1 Interaction domain |
| *MLH1* | rs63750449 | K618T | None/ S,P,R,M | 0.57% (NFE) | Y2H (8/8) | Exo1 Interaction domain |
| *MLH1* | rs3550531 | K618A | None/S,P | 0.34%** (overall) | ND | Exo1 Interaction domain |
| *MLH1* | rs63751049 | V326A | P*/S,R,M | 0.11% (SAS) | Y2H (4/5) | Transducer domain |
| *MLH1* | rs35831931 | V716M | None/ S,P,R,M | 0.20% (NFE) | Y2H(5/8) | C-terminal |

### Mouse modeling of nsSNPs and reproductive phenotypes

Based on the bioinformatic, biochemical and genetic data described above, we initially modeled five *MLH1* and three *MLH3* nsSNPs in mice via CRISPR/Cas9-mediated genome-editing (Table 1, Fig. S2). These eight mutant mouse lines with ‘humanized’ alleles were phenotyped in several ways, beginning with male reproductive parameters. Two mutants had obvious phenotypes. *Mlh3^R1230H/R1230H^* males exhibited a ~4 fold reduction in testes size, an absence of epididymal sperm, and histological evidence of meiotic prophase I arrest (Fig. 1 B-D). The phenotypes of *Mlh1^K618T/K618T^* males were less severe; their testes weighed 80% of WT, and there were ~35% fewer epididymal sperm (Fig. 1B-C). Fertility trials of homozygous mutant males confirmed that *Mlh3^R1230H/R1230H^* males were completely sterile, while *Mlh1^K618T/K618T^* males were subfertile, producing 40% fewer pups over a duration 10 months of fertility trials (Fig. 1H). In contrast to *Mlh1^K618T^* and *Mlh3^R1230H^*, there were no obvious reproductive defects in males homozygous for the remaining 6 alleles that were modeled. Sperm counts were normal and litter sizes sired by these mutant males were not lower than controls (Fig. 1C,H). Testis weights were also normal, though modestly lower in *Mlh3^R93G/R93G^* homozygotes.

Despite the absence of reproductive defects in the remaining 6 mutant alleles, testis sections revealed abnormal pyknotic-appearing metaphase spermatocytes with disorganized metaphase plates and misaligned chromosomes (Fig. 1D; explored in more detail below). This was more directly assessed by TUNEL staining of seminiferous tubule cross-sections positive for the metaphase marker phospho-histone H3 (Ser10) ^37^ which showed marked increases in apoptotic cells compared to WT (Fig. 1E-G). Confocal analysis readily revealed misaligned chromosomes at the metaphase plates of MI spermatocytes in the less severe mutants (Fig. S3). These data suggested that the allelic variants caused severe-to-subtle defects in chromosome alignment at the metaphase plate at MI, and/or unrepaired recombination intermediates, leading to apoptosis.

When this work began, the two K618 variants in *MLH1* (K618E and K618T) were (and are still) present in dbSNP as rs35001569 and rs63750449, respectively. These amino acid changes are caused by AAG><u>G</u>AG and AAG>A<u>C</u>G nucleotide changes, respectively. However, after those mouse models were created and phenotype analyses were performed (including female data presented below), the gnomAD database (v2.1) was revised to indicate that these two variants co-occur (are in phase) nearly perfectly (963/973 and 963/971 occurrences, respectively), encoding a K618A AA change as a result. This explains why both variants are annotated to occur at essentially identical frequencies in populations (Table 1). We therefore generated an *Mlh1^K618A^* allele. Testis histology appeared normal, and sperm counts between homozygotes and heterozygotes were similar implying that this allele is a neutral variant (Fig. S4).

### Meiotic recombination defects in mutants

To determine if defects in crossing-over might underlie the metaphase I aberrations, we immunolabeled pachytene spermatocyte chromosome spreads to quantify MLH1 foci, a proxy for “Type 1” crossovers (COs), which are subject to interference and constitute ~90% of all COs ^38^. In the two most severe mutants *Mlh3^R1230H/R1230H^* and *Mlh1^K618T/K618T^*, chromosomes exhibited normal homolog pairing and synapsis as reported for knockouts ^11,14,21^, but had markedly reduced MLH1 foci (Fig. 2A, B; 2.9 and 7.7 per cell, respectively, *vs*. 21.4 foci in WT). Such a severe reduction in ‘obligate’ class I COs might either trigger apoptosis at metaphase I, or prevent zygotic progression after fertilization ^21,26^.

**Figure 2.**
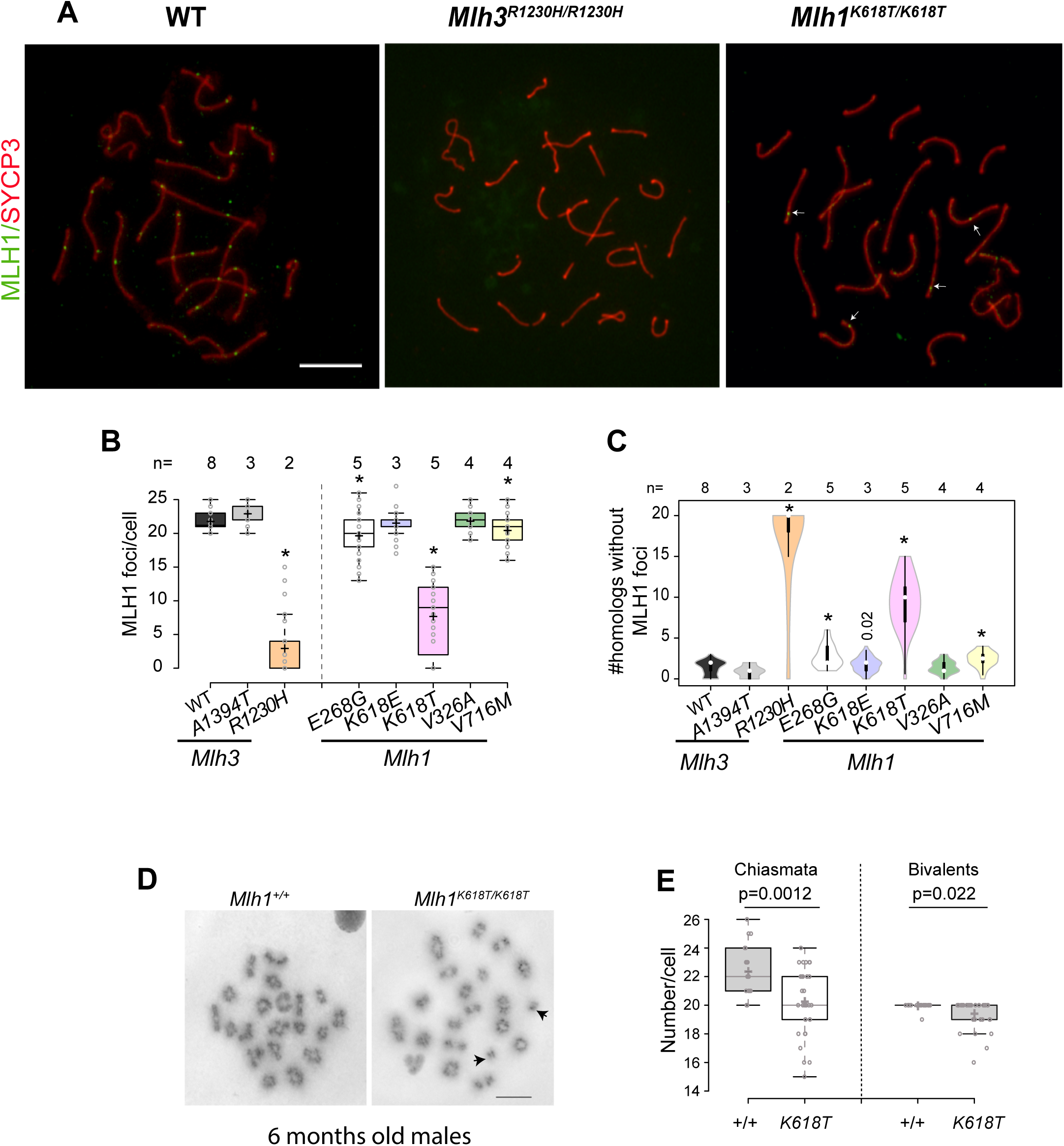
Recombination in mutant spermatocytes. **A)** MLH1 immunolocalization in spermatocyte nuclei. Shown are pachytene spermatocyte chromosome surface spreads from 6 month old males, immunolabeled as indicated. White arrows indicate MLH1. Note weak and nearly absent staining in the mutants. **B)** MLH1 focus quantification in spermatocytes. * p value <0.0001. Others do not have p values <0.05 for decreased foci. Over 45 cells from at least 3 animals were quantified, except for *Mlh3^R1230H^*, where 26 cells from two animals were scored, and *Mlh3^A1394T^*, where 29 cells from three animals were scored. **C)** Violin plots illustrating the chromosome pairs in a spermatocyte nucleus lacking an MLH1 focus. White circles show the medians; box limits indicate the 25th and 75th percentiles; whiskers extend 1.5 times the interquartile range from the 25th and 75th percentiles; polygons represent density estimates of data and extend to extreme values. For mutants with more MLH-deficient pairs, * = p value <0.0001 unless otherwise indicated; comparisons to WT with p values >0.05 are not shown. **D)** Example of spermatocyte metaphase spread from the *Mlh1^K618T^* mutant and control. **(E)** Quantification of chiasmata and bivalents in *Mlh1^K618T^* and WT. Twenty or more cells from three animals were analyzed. Scale Bar = 5μm. Data in B and E are plotted as follows: Boxes indicate data points between the first and third quartiles of the distribution, whiskers show minimum and maximum values, black horizontal lines and ‘+’ sign indicate the median and mean values respectively.

The *Mlh1^V716M^* and *Mlh1^E268G^* mutants, though less affected, also had significantly fewer total MLH1 foci per spermatocyte on average. Since even small reductions in CO recombination might lead to some chromosomes lacking an obligate CO (chiasma), we quantified MLH1 focus-deficient chromosomes. Five of the seven mutants examined exhibited a significant increase in MLH1 focus-deficient chromosomes (Fig. 2C). This suggests an increase in spermatocytes bearing one or more achiasmate chromosomes. Consistent with this hypothesis, *Mlh1^K618T/K618T^* spermatocytes exhibited an average of ~2 fold fewer metaphase I chiasmata than controls, leading to a significant decrease in bivalent chromosomes (increase in univalents; Fig. 2D).

### Age-dependent decline in reproductive capacity of *Mlh1/3* mutant females associated with increased oocyte aneuploidy

In *Mlh1* or *Mlh3* knockouts, females are sterile because their oocytes cannot form a normal spindle and chromosomes do not properly align to the meiotic metaphase I plate ^30,39^. This near complete absence of bi-oriented, CO-tethered bivalents at diakinesis can activate quality control mechanisms, namely the Spindle Assembly Checkpoint (SAC), causing meiotic arrest ^26,28^. The SAC appears to be more stringent in spermatocytes than oocytes ^27,29,40,41^, thus eliminating defective spermatocytes whereas oocytes attempt to complete gametogenesis. To test potential impacts on fertility of females bearing the humanized *Mlh1/3* alleles, we evaluated fecundity of the *Mlh1^E268G^, Mlh1^V326A^, Mlh1^K618E^, Mlh1^K618T^*, and *Mlh1^V716M^* mutants. Interestingly, litter sizes of all mutant females declined over time compared to controls, and overall, had only ~ half of the total offspring as controls over the 36 weeks they were housed with WT sires (Fig. 3A, B). The overall shortfall during this period was largely attributable to: **i)** decreases in fecundity after the first four litters, when most dams were ~5-5.5 months of age (Fig. 3a), and **ii)** ceasing to have litters at an earlier age than WT, despite evidence for continued ovulation (Fig. S5). Whereas homozygotes of the aforementioned five alleles showed age-related decrease in litter sizes, only *Mlh1^K618T/K618T^* females had dramatically fewer (~half) follicles (combined primordial and developing follicles, Fig. 3C) at three weeks of age. While we did not perform longitudinal studies on the *Mlh1^K618A^* allele, female homozygotes were fertile. In preliminary crosses, two young homozygous females had apparently normal litter sizes – an average of 7.5 pups/litter (N=5 litters) – when crossed to heterozygous males.

**Figure 3.**
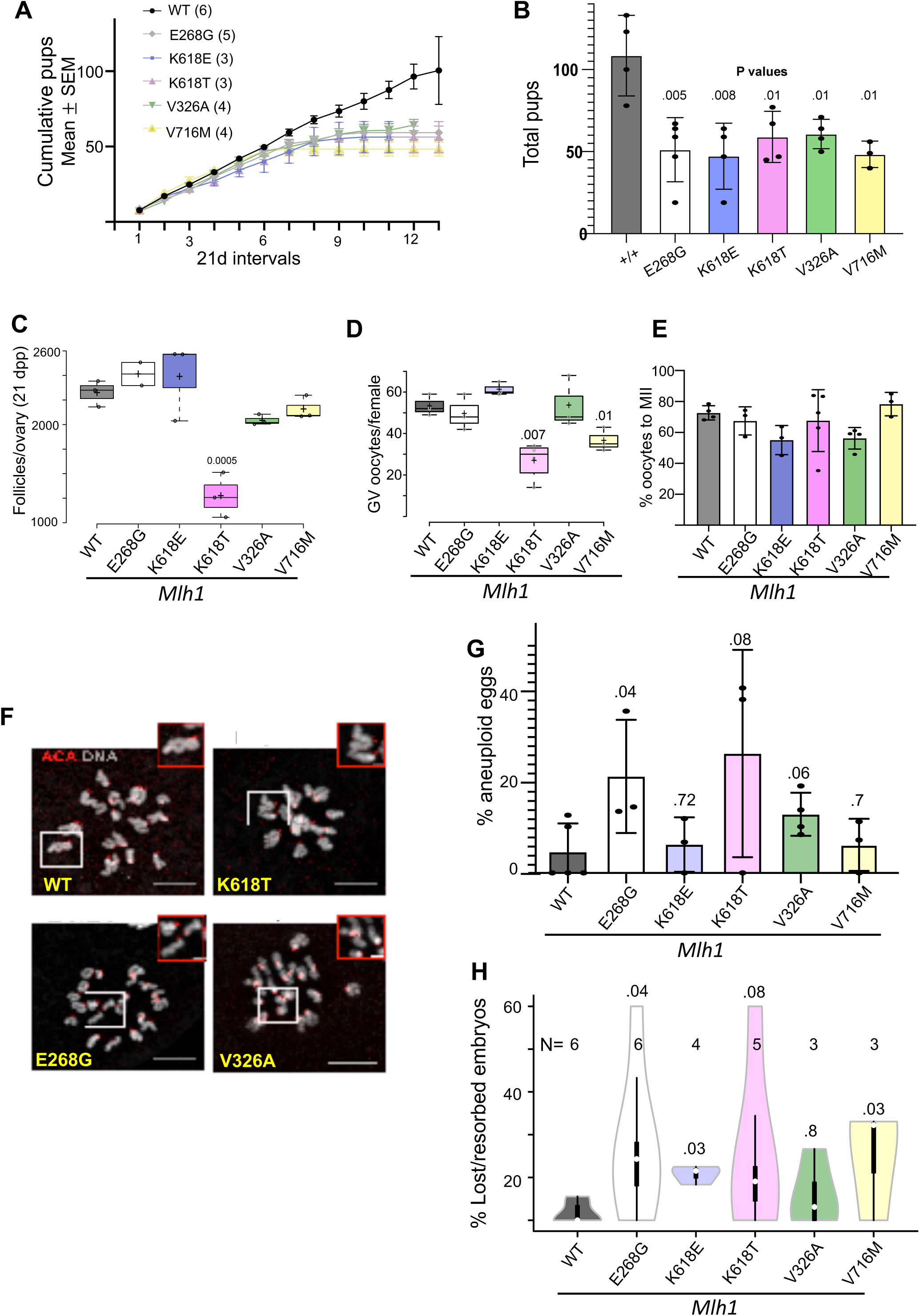
Age-dependent subfertility in *Mlh1* mutant females and increased increased aneuploidy in oocytes. **A)** Litter sizes of mutant females over their reproductive lifespans. Eight-week old wild-type and littermate mutant females were mated with wild-type males, and the number of pups born per litter are plotted. For all panels, the following abbreviations apply to genotypes: E268G, *Mlh3^E268G/E268G^*; K618E, *Mlh1^K618E/K618E^*; K618T, *Mlh1^K618T/K618T^*; V326A, *Mlh1^V326A/V326A^*; V716M, *Mlh1^V716M/V716M^*. **(B)** Total pups delivered by females during their reproductive lifespan (~2 to 10 months, see Methods). **(C)** Follicle counts (all stages; the great majority were primordial follicles) in three-week old females. P values <0.05, in comparisons of mutant to WT, are indicated. Dpp, days post-partum. **(D)** Number of total germinal vesicle (GV) stage oocytes collected from superovulated WT and mutant females (3-4 month old). p values <0.05 relative to WT (+/+) are shown. Horizontal lines are mean and whiskers are ± Std. dev. **(E)** Percent maturation of GV oocytes (shown in A) matured to metaphase meiosis II (Met II). p values <0.05 relative to WT (+/+) are shown. **(F)** *In situ* metaphase chromosome spreads showing abnormalities in mutants. Representative confocal z projections are shown. Oocytes were stained to detect centromeres (ACA, red) and DNA (DAPI, gray). Scale Bar = 10 μm in main panels, and 2μm in insets; F= 20 mm. **(G)** Percentage of aneuploid metaphase II oocytes from (E). p values are shown above bars. **(H)** Violin plots of embryo loss estimates in *Mlh1* mutants. Females homozygous for the indicated mutations (3-4 mth old) were mated to C57BL/6J (WT) males, sacrificed at E13.5, and the number of viable embryos scored, as well as ovulation sites on the ovary. The difference between the two were presumed to have died during gestation at various stages, and are plotted on the Y axis. The control females (“WT”) were +/+ or heterozygous littermates. p values are shown, and were calculated from the Student’s two-tailed t-test. Whiskers extent represent minimum and maximum values. N values are number of pregnancies examined.

It is possible that in the *Mlh1^K618T^* mutants with reduced follicles at three weeks of age (and which also had the greatest reduction in COs in males), oocyte loss was triggered by failure to repair CO-designated DSBs, and subsequent activation of the meiotic DNA damage checkpoint ^42^. Alternatively, the smaller litter sizes from mutant dams might be attributable decreased COs and thus increased aneuploidy in ovulated eggs with age, although it is unclear how this would occur. An increase in aneuploidy could lead to increased embryonic lethality as with *Sycp3* null females ^9^.

To test the hypothesis that reduced litter sizes are attributable to aneuploidy-induced embryo loss, we first examined the fraction of aneuploid eggs produced by mutant females (3-4 months of age). Following hormone stimulation, oocytes were matured *in vitro* to MII, then spread to visualize chromosomes. All mutants examined except *Mlh1^K618T^* and *Mlh1^V716M^*, both of which had lower ovarian reserves (see above), contained similar numbers of prophase I arrested oocytes as WT females (Fig. 3D). About 70% of the oocytes from all genotypes matured to MII after 15 hrs (Fig. 3E), although maturation, as marked by polar body extrusion, was ~20% less efficient for the *Mlh1^K618E^* and *Mlh1^V326A^* genotypes. The *Mlh1^E268G^*, *Mlh1^K618T^* and *Mlh1^V326A^* alleles caused marked increases in aneuploid eggs, detected as abnormal numbers of kinetochores/chromatids at MII (Fig. 3F, G; 20.9±7.1%, 25.8±12.9% and 12.7±2.3, respectively, compared to 4.5±2.7% in WT).

We next explored whether the high levels of aneuploidy in eggs from mutant females led to increased embryonic death of embryos. Control and mutant females that were 3-4 months old (similar to the age of females used in the ploidy experiments) were mated with wild-type males, dissected at E13.5, and the numbers of viable embryos and corpora lutea (CL; reflecting the number of ovulated oocytes) were counted. The fraction of embryonic lethality was inferred as: [CL-embryos]/CL ^43^. Whereas lethality was only 3.4% in WT mothers, it was several fold higher in the mutants, ranging from 11.4% in *Mlh1^V326A^* to 34.5% in *Mlh1^E268G^* (Table 2). These results suggest that mutations in MLH1/3 can elevate the incidence of aneuploid oocytes, leading to a corresponding increase in spontaneous pregnancy loss.

**Table 2.**
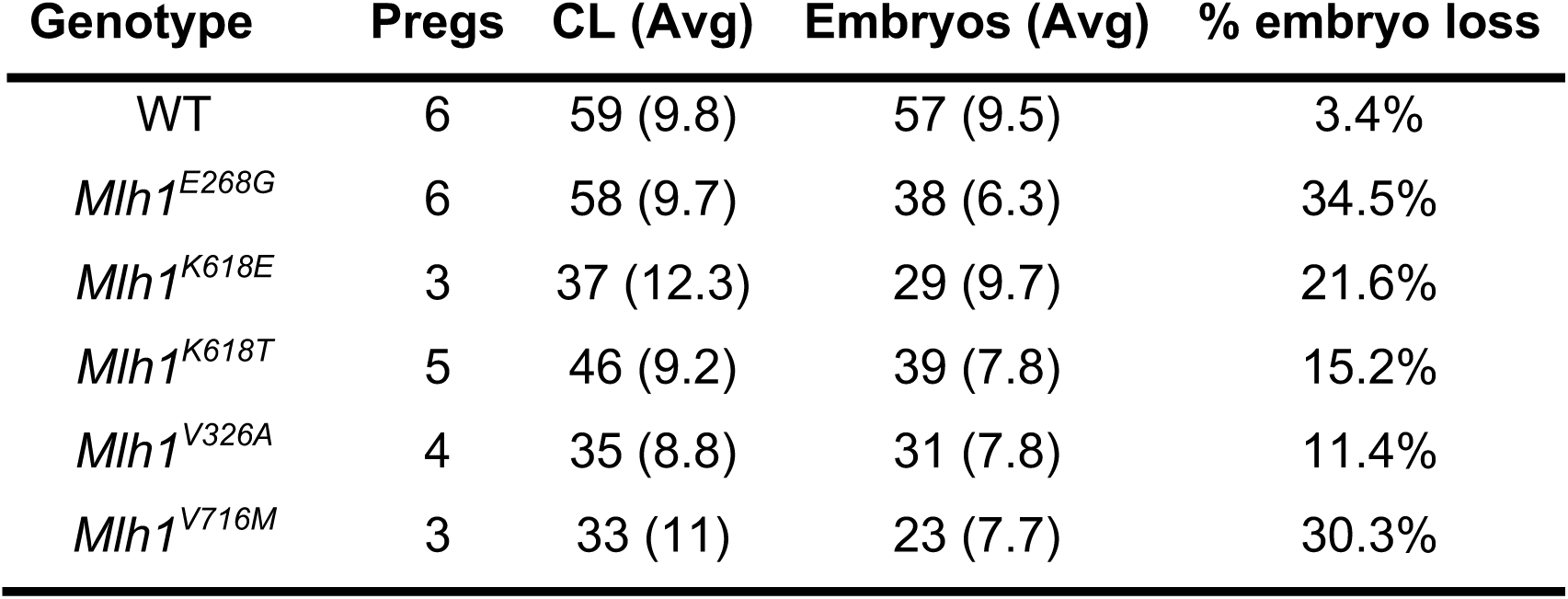
Embryo loss during development. All mutant genotypes are homozygous. # Pregs, pregnancies dissected; CL, Corpora lutea; Avg, average. The embryos counted were viable when dissected at E13.5. The % embryo loss = (#CL - #embryos)/#CL * 100.

## Discussion

The process of meiotic recombination, specifically crossing over, is essential for fertility and the production of viable, healthy offspring. A severe reduction in COs will prevent proper alignment of meiotic chromosomes at the metaphase I plate, leading to arrest or failed fertilization. However, if only one or a very few chromosomes lack a chiasma, viable gametes can still form but will be prone to mis-segregation of parental homologs; this can lead to aneuploid embryos. A recent study of recombination in human fetal oocytes revealed that having 10-20% fewer COs (MLH1 foci) overall was associated with such oocytes having 1-2 chromosomes completely lacking a CO ^44^. Given that the most common cause of miscarriage (~60%) is embryonic aneuploidy, primarily trisomies ^1^, it is of interest whether the frequency of so-called “exchangeless” chromosomes is genetically controlled. There are indications that parental genetic factors can predispose to generation of aneuploid conceptuses, and may contribute to cases of RPL ^5,6^. In this respect, variants impacting genes important for recombination, including the MUTL homologs, are potential candidates for influencing aneuploidy rates.

SNPs in *MLH1* and *MLH3* have been statistically associated with male infertility in some studies focusing on candidate genes (i.e., not GWAS), but there haven’t been validation studies ^45–47^. For female fertility, GWAS for phenotypes such as age to menopause, polycystic ovary syndrome, and endometriosis have been performed, but only a few associations have been reported for genes involved in oogenesis *per se* ^48^. In this study, we focused on human Variants of Unknown Significance (VUS) in MLH1/3, with the goal of determining possible effects on fertility. Because our earlier studies of VUS in essential fertility genes indicated that functional prediction algorithms alone overpredicted deleteriousness ^49^, we applied additional functional criteria to select variants for mouse modeling, namely, studies of analogous mutations in yeast or disruption of protein-protein interactions in Y2H assays. Given that most of the alleles reported here that exhibited defects in these assays caused phenotypes in mice, we believe that this approach is valuable for prioritizing experimentation.

In our opinion, the most intriguing result from this study is that female fecundity in several mutants declined with age and reproduction ceased prematurely, despite having no apparent shortfall (compared to WT) in the ovarian reserve at wean age. One possibility is that after wean age (~3 weeks), oocyte attrition accelerated in the mutants, such that they exhausted their ovarian reserve. However, histology of these older females revealed numerous corpora lutea, indicative of continued ovulation in these aged females (Fig. S5). Future experiments aimed at quantifying the reserve, numbers of ovulated eggs, the ability of these eggs to become productively fertilized, and other markers of ovarian aging could reveal the basis of this phenomenon.

An alternative explanation is that the quality of oocytes recruited later in life might be lower than at earlier stages in the mutants. We can only speculate as to the nature of such quality differences, but they might be related to increased aneuploidy caused by prolonged periods in primordial follicles with achiasmate chromosomes. It is also possible that these mice might represent an example of the “production line” hypothesis that was originally proposed as an explanation for increased aneuploidy in women as they age ^50^. This hypothesis posits that eggs are ovulated in the order that they originally proceed through meiosis *in utero*, and that the “later” oocytes suffer from deficient recombination, and thus higher aneuploidy at MI. While this hypothesis has been refuted in humans ^51^, there is evidence that in mice, the first oogonia to enter meiosis are the first to be ovulated ^52^. Nevertheless, there is no evidence relating crossover rates to time of meiotic entry. Interestingly, a similar phenomenon was observed in *Sycp3* null mice, where an age-correlated decrease in litter sizes was attributable to increased embryonic death from aneuploidy ^9^. Additionally, a small scale study implicated mutations in *SYCP3* as being responsible for RPL in two women ^7^. It has been hypothesized that women with smaller ovarian reserves have an increased chance of having aneuploid abortuses ^53^. In future work, our models may be useful in exploring this possible relationship in aging mice, and to better understand sexually dimorphic consequences upon gamete quality and checkpoint mechanisms.

Inherited mutations in MLH1 can cause Lynch Syndrome, sometimes referred to as hereditary non-polyposis colorectal cancer (HNPCC), although patients are also subject to a variety of cancers including endometrial ^54^. Lynch Syndrome exhibits a dominant mode of inheritance, and the tumors are characterized by microsatellite instability (MSI). Tumor initiation is attributable to either epigenetic silencing or mutation of the normal *MLH1* allele inherited from one parent ^54–56^. Mice null for *Mlh1* exhibit MSI, increased morbidity (moribund at an average age of 7 months), and develop lymphoma and various types of tumors including gastrointestinal ^57^. However, we did not observe an increased incidence of tumors in any lines or microsatellite instability in the mutants tested (data not shown). Mutations in *MLH3* are not associated with Lynch Syndrome, but there is one report that null mice are predisposed (50%) to gastrointestinal tumors with a long latency ^58^. Although the orthologous yeast *mlh3* alleles have MMR defects (Table 1), we did not observe tumors or premature morbidity in our *Mlh3* mouse mutants. However, we did not age them beyond 1 year. It is also possible that for both sets of models (*Mlh1/3*), that the genetic background we used was not as tolerant to transformation (at least before 1 year of age) as those in published studies. Because of these caveats, we cannot conclude that these alleles do not cause Lynch Syndrome in humans.

As mentioned in the Results, when this project began, we selected the two variants at K618 (K618E and K618T) that remain present in dbSNP as rs35001569 and rs63750449, respectively. However, after the mouse models were created, the gnomAD database was revised to indicate that these two variants co-occur (are in phase), encoding a K618A amino acid change at essentially identical frequencies in populations (Table 1). The question remains as to whether the *MLH1^K618E^* and *MLH1^K618T^* exist at all in populations, or at very low frequencies (approximately 100-fold lower based on the numbers cited above). Notably, the *MLH1^K618E^* and *MLH1^K618T^* alleles are listed in the Clinvar database, and reported in patient samples ^59–61^. It is unclear if the three alleles arose independently, or if they were derived from one another. Regarding the *MLH1^K618A^* allele and its relationship to Lynch Syndrome, which is somewhat controversial, a recent large study found no evidence for this allele’s involvement, and thus concluded that *MLH1^K618A^* is benign ^62^. Interestingly, a computational and experimental study of *MLH1* variants suggested that many variants that cause Lynch Syndrome do so by thermodynamically de-stabilizing the MLH1 protein, leading to increased degradation ^63^. Abildgaard and colleagues tested four of the alleles modeled here (*MLH1^E268G, K618T, K618A and V716M^*), and each were classified as being “likely benign” by virtue of the proteins having substantially normal stability.

In sum, these studies indicate that variants in the MUTLg (MLH1/3) complex genes can have subtle-to-severe effects on gametogenesis that may go unnoticed without rather detailed phenotyping, including aging studies. Homozygosity for most of the alleles did not cause infertility or drastic reductions in ovarian reserve or sperm counts, although it is conceivable that some of these alleles in *trans* to a null allele, or in the context of a variant in the other member of the MUTLg complex, could cause synthetic phenotypes. Nevertheless, the phenotypes of increased aneuploidy in oocytes, and reduced reproductive lifespan of females are highly relevant to the human condition, and thus these mouse models can be valuable for assessing risk at an early age in women, and to dissect the biology of these phenomena.

## MATERIALS AND METHODS

### Construction of baker’s yeast strains to measure meiotic crossing over and DNA mismatch repair

The SK1 background baker’s yeast strain EAY3255 was constructed for the analysis of MMR and meiotic crossing over phenotypes ^64^. These and all yeast strains generated/used are listed in Table S4. It contains a spore autonomous fluorescent protein (Tomato) marker located near the centromere on chromosome VIII ^36^ and a *lys2::InsE-A14* cassette to measure reversion to Lys^+ 35^. *mlh3* alleles cloned into pEAI plasmids (*mlh3-X::KANMX*) were introduced by gene replacement into EAY3255 using the lithium acetate transformation method ^65^. These plasmids were digested with *Bam*HI and *Sal*I prior to transformation. At least two independent transformants for each genotype (verified by DNA sequencing) were made. EAY3255 and derivatives were used to measure the effect of *mlh3* mutations on *lys2::InsE-A14* reversion rate. To measure meiotic crossing over, these strains were each mated to EAY3486, an SK1 *mlh3Δ* strain containing the m-Cerulean fluorescence marker located at *THR1* on chromosome VIII, ~20 cM away from the Tomato marker present at *CEN8* in EAY3255. Diploids were selected on media lacking the appropriate nutrients and maintained as stable strains. Meiosis was induced upon growing the diploid strains on sporulation media as described in ^66^. Wild-type strains carrying the fluorescent protein markers used to make the above test strains have been described ^36^.

### Spore autonomous fluorescent protein expression assay to measure meiotic crossing over

Tetrads obtained from EAY3255/EAY3486 background diploids were treated with 0.5% NP40 and briefly sonicated before analysis by fluorescence microscopy. Tetrads were analyzed using a Zeiss AxioImager M2 microscope equipped with RFP and CFP filters. At least 250 tetrads for each *mlh3* allele were counted to determine the % tetratype (two spores show the parental genotype and two show a configuration consistent with a crossover event between the Tomato and m-Cerulean markers located on Chromosome VIII; Table S3). Two independent transformants were measured per allele, measured on at least two days with a similar amount analyzed to avoid batch effects. A statistically significant difference (p<0.002) from wild-type and *mlh3Δ* controls, based on χ^2^ analysis (Pearson χ^2^ contingency test, with a Bonferonni correction for 26 comparisons), was used to classify each allele as exhibiting a wild-type (+), intermediate (+/-, +/--), or null (-) phenotype.

### *lys2_A14_* reversion assay to monitor MMR

The EAY3255 derived haploid strains described above were analyzed for reversion to Lys+ as described ^35^. 11 to 16 independent cultures were analyzed for each mutant allele as well as wild-type and *mlh3Δ* controls. Two independently constructed transformants for each allele were analyzed on at least two days to avoid batch effects, with a similar number of repetitions performed each day. Reversion rates (Table S3) were measured and the median rate for each genotype was normalized to the wild-type median rate (1x) to calculate fold increase. Alleles were classified as wild-type (+), intermediate (+/-), or null (-) based on the 95% confidence intervals (CI) ^67^. If the 95% CI for the allele overlapped with the 95% CI for one of the controls (wild-type or null) they were considered statistically equivalent to the respective control.

### Variant selection for *MLH1*

To select *MLH1* SNPs for cloning and subsequent Y2H testing, we filtered gnomAD v2.1 for missense mutations with overall AF between 0.02% and 2%. 15 missense SNPs fell within this range, of which 13 were successfully cloned and tested by Y2H.

### Generating mutant *MLH1* clones

Clones for *MLH1* and all Y2H-tested interaction partners were obtained from hORFeome v8.1 (Yang et al., 2011) or v5.1 (The MGC Project Team, 2009). The *MLH1* alleles were generated by site-directed mutagenesis. Primers for mutagenesis were designed using the webtool primer.yulab.org. To minimize sequencing artifacts, PCR was limited to 18 cycles using Phusion polymerase (New England Biolabs, M0530). Mutagenesis PCR product was then digested overnight using *DpnI* (New England Biolabs, R0176) followed by bacterial transformation into competent cells. Single colonies were recovered on LB agar plates containing spectinomycin. Four colonies were picked per mutagenesis attempt and then were validated to contain to the mutation of interest through Sanger sequencing using primers designed to cover the entire *MLH1* sequence.

### Profiling disrupted protein interactions through Y2H

Wild-type and successfully mutated *MLH1* clones were transferred into Y2H vectors pDEST-AD and pDEST-DB by Gateway LR reactions then transformed into *MAT***a** Y8800 and *MAT*α Y8930, respectively. Testable Y2H interaction partners were identified by surveying a Y2H reference interactome comprised of interactions reported in four manuscripts (Rolland et al., 2014; Rual et al., 2005; Venkatesan et al., 2009; Yu et al., 2008). All ORFs corresponding to interactions with MLH1 were then selected for Y2H testing. MLH1 interaction partners, transformed into *MAT***a** Y8800 and *MAT*α Y8930 were then mated against AD- and DB-MLH1 transformed yeast of the opposite mating type in a pairwise orientation by co-spotting yeast inoculants onto YEPD agar plates. To screen for autoactivators, DB-MLH1 yeast cultures were also mated against *MAT***a** yeast transformed with empty pDEST-AD vector. Mated yeast were incubated overnight at 30°C and then replica-plated onto dropout Synthetic Complete agar media lacking leucine and tryptophan (SC-Leu-Trp) to select diploid yeast. After overnight incubation at 30°C, diploid yeast were then replicate-plated onto dropout SC agar media lacking leucine, tryptophan, histidine, and supplemented with 1 mM of 3-amino-1,2,4-triazole (SC-Leu-Trp-His+3AT). Diploid yeast were also replica-plated onto SC agar media lacking leucine, tryptophan, and adenine (SC-Leu-Trp-Ade). After overnight incubation at 30°C, plates were replica-cleaned and incubated again for three days at 30°C. An interaction was scored as disruptive only if (1) mutant MLH1 reduces growth by at least 50% relative to its corresponding wild-type interaction, and (2) neither wild-type nor mutant DB-MLH1 scored as an autoactivator.

### Production of ‘humanized’ mouse-lines and mouse breeding

All ‘humanized’ alleles were generated by CRISPR/Cas9-mediated genome editing as we have described ^49,68^. Briefly, Cas9 protein, sgRNA and single stranded oligodeoxynucleotides (ssODN) bearing the desired sequence changes (oligos listed in Table S5) were microinjected into single cell zygotes produced from matings between FVB/NJ and B6(Cg)-Tyr^c-2J^/J) inbred mice. Correctly edited founder (F0) animals were backcrossed to B6(Cg)-Tyr^c-2J^/J for two generations, then maintained in a mixed strain background (C57BL/6J x FVB/NJ).

### Fertility tests and embryo loss

Control and experimental animals (8-10 weeks old) were housed with age-matched fertile mates (C57BL6/J) for up to 10-12 months of age. Litter sizes were determined by counting pups on the day of birth.

For embryo loss analyses, homozygous females (12-16 weeks old) were mated to WT males and the presence of a copulation plug in the morning was recorded as 0.5 dpc. Females were sacrificed at 13.5 dpc for counting of both implantation sites and corpora lutea (CL; indicating the numbers of ovulated oocytes). Graphs and statistical analyses were performed with GraphPad Prism5.

### Ovary histology and follicle quantification

Ovaries were collected from 3-week females and were fixed in Bouin’s for 24h, washed in 70% ethanol, paraffin embedded, and further serial sectioned at 6 μm followed by staining with hematoxylin and eosin (H&E). Every fourth section was counted. Though we reported the combined number of follicles in Figure 3, we maintained records of the follicle subtypes (e.g., primordial, primary, secondary, antral) and can provide them upon request.

### Surface spread preparation and immunocytology

Prophase I surface spreads were prepared as described ^69^. For all experiments, at least 3 males from each genotype was evaluated. Following final washing of chromosome slides in 0.4% Kodak Photo-Flo 200/dH2O for 2 x 5 min, slides were air dried for ~10 min and stored in −80°C or used immediately for staining. Primary antibodies used were: rabbit anti-SYCP3 (1:500, ab15093; Abcam), mouse anti-SYCP3 (1:500, ab97672; Abcam), and mouse anti-MLH1 (1:50, 550838; BD Pharmingen). Secondary antibodies were goat anti rabbit-IgG 488 (1:2,000, A11008; Molecular Probes), goat ant-rabbit IgG 594 (1:1,000, A11012; Molecular Probes), and goat anti-mouse IgG 594 (1:1,000, A11005; Molecular Probes). Nuclei were counterstained with DAPI, and slides were mounted with ProLong Gold antifade reagent (P36930; Molecular Probes).

### Sperm counting

This was performed as described previously ^49^.

### Testes histology and Immunohistochemistry

For histological analyses, testes were fixed for 24 h at room temperature (RT) in Bouin’s solution. Tissue was further paraffin-embedded, sectioned (7 μm), and stained with H&E. For immunohistology, testes were essentially fixed in 4% paraformaldehyde for ~24 h. Paraffin-embedded tissues were further sectioned at 7 μm. For TUNEL and PH3 double immunostaining, sections were deparaffinized followed by antigen retrieval using Sodium Citrate Buffer (10mM Sodium Citrate, 0.05% Tween 20, pH 6.0). Sections were blocked in PBS containing 5% goat serum for 1 h at RT, followed by TUNEL staining, performed following instructions manufacturer’s instruction (Invitrogen, Click-iT™ TUNEL kit). This was followed by 1° antibody incubation overnight at 4°C (anti-pH3 (Ser10), 1:100, Millipore) and anti-rabbit AF488 (Invitrogen, 1:1000) 2° antibody incubation for 1 h, RT. Slides were counterstained using DAPI, mounted in Vectashield and imaged as described in below.

### Imaging and analysis of testes cross sections

Testes were fixed in Bouin’s, embedded in paraffin and stained with hematoxylin and eosin (H&E) in Cornell’s Diagnostic Pathology Lab. For TUNEL labeling, testes cross sections were double immunolabeled with TUNEL and anti-pH3 (see above) antibody and were imaged using a confocal microscope (u880, Carl Zeiss, Germany) with a Plan Apo 40× water immersion objective (1.1 NA) and Zen black software. The following lasers were used: argon laser-488 nm, blue-diode-405 nm, DPSS laser—561 nm. Following identical background adjustments for all images, cropping, color, and contrast adjustments were made with Adobe Photoshop CC 2017.

### Oocyte *in vitro* maturation and *in situ* chromosome counting

Prophase I-arrested oocytes (GV) were collected as described ^70^. Forty-eight hours prior to collection, females were injected intraperitoneally with CARD HyperOva (KYD-010-EX-X5; CosmoBio USA) consisting of Inhibin antiserum and equine chorionic gonadotrophin. Briefly, ovaries from 12-16 weeks old females were dissected and GV oocytes were released from the follicles by puncturing the ovaries several times with 27-gauge sewing needles in MEM/polyvinylpyrrolidone media containing 2.5 μM milrinone (Sigma-Aldrich; M4659) to prevent oocyte maturation after the release from cumulus cells. Then, cumulus cells were removed by gentle pipetting and fully-grown oocytes were matured in vitro in Chatot, Ziomek, and Bavister (CZB) media without milrinone in a humified incubator programmed to 5% CO_2_ and 37°C for 14h until they arrived to metaphase II. They were cultured for 2h in 100 μM Monastrol (Sigma, M8515) to disorganize the spindle and separate the chromosomes. Then, eggs were fixed in 2% paraformaldehyde in PBS for 20 min and permeabilized in PBS containing 0.2% Triton X-100 (Sigma-Aldrich, 900-93-1) for 20 min. Eggs were stained with the anticentromere antibody (ACA) to detect centromeres (Antibodies Incorporated; #15-234; 1:30) and DAPI to detect DNA. Chromosome numbers were quantified in metaphase II eggs as described ^71^. Mouse eggs should have 20 pairs of sister chromatids; any deviation of this number was considered aneuploid. Eggs were imaged with the i880 (Zeiss) confocal at 0.5 μm z-intervals. Chromosome counting was performed with NIH Image J software, using cell counter plugins.

## Acknowledgements

We thank S. Keeney for yeast strains, R. Munroe and C. Abratte in Cornell’s transgenic core for making mice. This work was supported by grants R01HD082568 to JS and HY, R01HD091331 to KS, and R35 GM134872 to EA.

