## Supplemental Figs 1-5 for "*MLH1/3* variants causing aneuploidy, pregnancy loss, and premature reproductive aging"

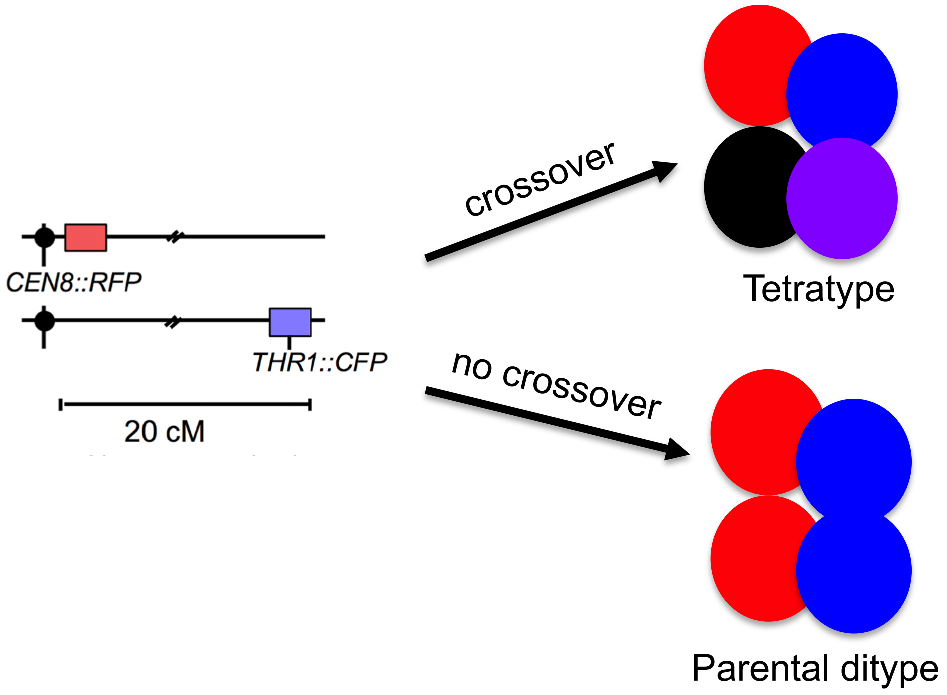

**Fig. S1. Spore autonomous fluorescence assay (Thacker et.al, 2011) to monitor crossover frequencies.** Diploid yeast bearing mutant *mlh3* alleles and indicated red (RFP) and cyan (CFP; colored blue in cartoon) fluorescence markers knocked into loci on Chr VIII are sporulated. Sister chromatids not shown. Segregation of single fluorescence-positive spores (blue or red) indicates no crossovers. Double positive (purple; lower right circle in the tetratype example) or fluorescence-negative spores (black circle) is indicative of a crossover.

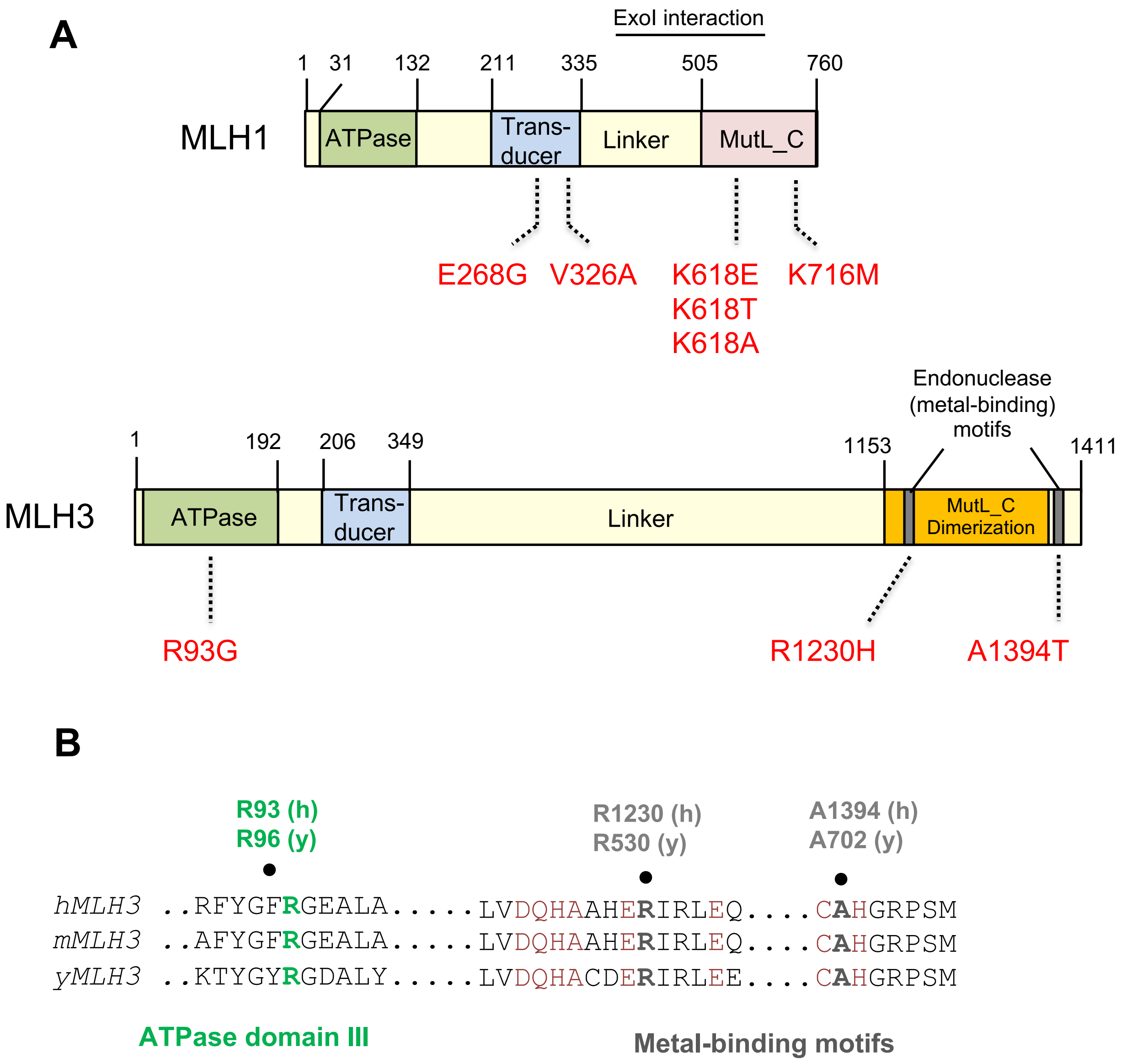

**Figure S2. Cartoon of mouse MLH1 and MLH3 protein organization, and locations of variants modeled in mice. A)** Amino acid (AA) positions in full-length isoforms, above the gene diagrams, correspond to the mouse proteins. The human variants modeled are noted in red below the protein diagrams, and those AA positions correspond the the human proteins. Indicated in the diagrams (colored boxes,except for the Exo[nuclease]I interaction domain) are approximate locations of domains in each protein. **B)** Locations (black dots) of AA alterations modeled in conserved ATPase and endonuclease domains in yeast and mouse MLH3. h = human; m = mouse; y = yeast. The key Aas for the metal binding motifs are in red, based on homology to residues in yeast Pms1 that form a metal binding site in the C-terminal domain of yeast Mlh1-Pms1 (Gueneau et al. 2013). Conserved ATPase domains are based on (Ban and Yang 1998; Tran and Liskay 2000).

• Gueneau E, Dherin C, Legrand P, Tellier-Lebegue C, Gilquin B, Bonnesoeur P, et al. (2013) Nat Struct Mol Biol 20: 46-468.
• Tran PT, Liskay RM (2000). Mol Cell Biol 20: 6390–6398.
• Ban C. Yang W (1998). Cell 95: 541–552.

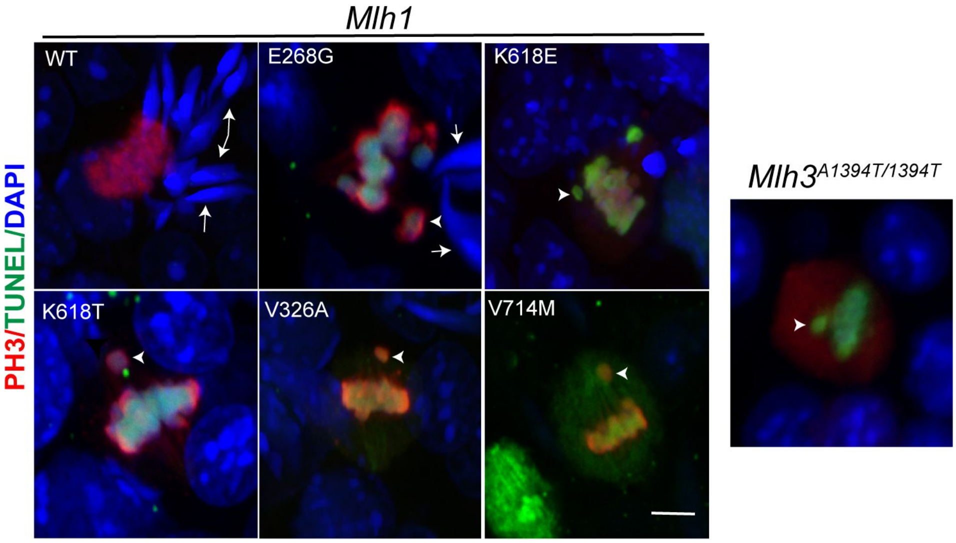

**Fig S3. Confocal projected images of MI spermatocytes illustrating misaligned chromosomes.** Testes cross sections stained with antibody against phosphorylated histone H3 (Ser10) (red) and TUNEL (green) to visualize apoptotic metaphase cells. Arrowheads point to misaligned chromosomes. Arrows indicate elongated spermatids. Scale bar, 5 um

## A

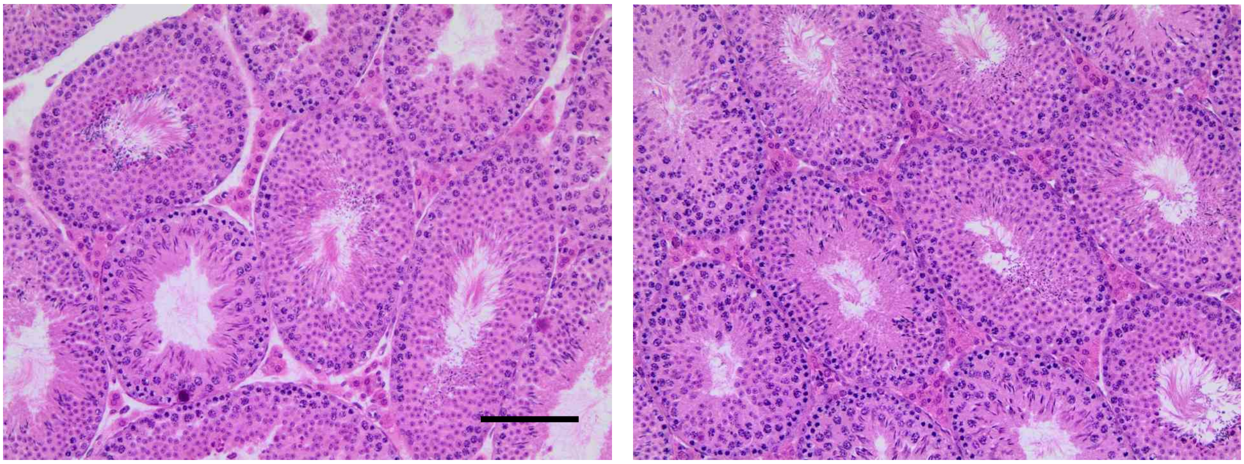

*Mlh1*$^{K618A/+}$ *Mlh1*$^{K618A/K618A}$

## B

| Genotype | Sperm/ml |
|---|---|
| *Mlh1*$^{K618A/K618A}$ | 5.3 x $10^5$ |
| *Mlh1*$^{K618A/+}$ | 4.6 x $10^5$ |
| *Mlh1*$^{K618A/+}$ | 4.6 x $10^5$ |
| *Mlh1*$^{K618A/K618A}$ | 5.7 x $10^5$ |
| *Mlh1*$^{K618A/+}$ | 5.1 x $10^5$ |
| *Mlh1*$^{K618A/+}$ | 4.9 x $10^5$ |

**Figure S4 – Testis histology and sperm counts of *Mlh1*$^{K618A}$ mutants.** A) Representative H&E-stained paraffin sections of testes of indicated genotypes. Mice were 87 days old. Scale bar = 50 µm. B) Sperm counts of 6 adult mice of the indicated genotypes. The concentration is of 2 ml of sperm obtained from a single minced cauda epididymis (see Methods).

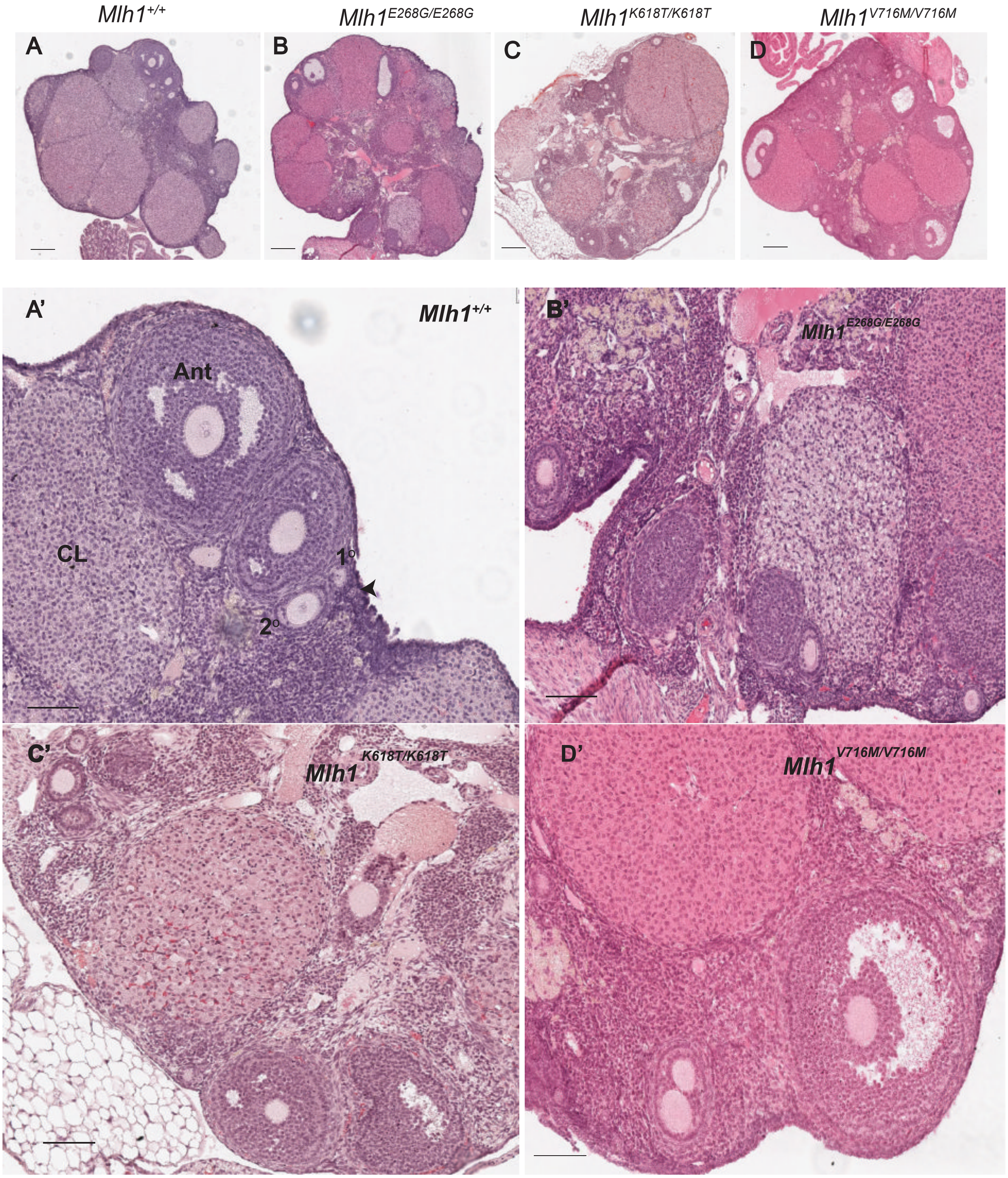

**Fig S5. *MLH1* SNPs do not abolish ovulation in aged WT and *Mlh1* mutant females.** H&E stained mouse ovaries from >11 month old females of indicated genotypes (abbreviation consistent with previous figures). Scale bar= 200μm in A-D and 100 μm in A'-D'. An- Antral follicle; 1º- primary follicle; 2º- secondary follicle; CL- Corpus Luteum; arrowhead- primordial follicle.
