## Supplementary Table 2 for "*MLH1/3* variants causing aneuploidy, pregnancy loss, and premature reproductive aging"

**Table S2 - Analysis summary of yeast equivalents to human *MLH3* SNPs for defects in crossing over and mismatch repair.**

| Variant | SNP ID | Yeast  Equiv | MMR | CO | Domain |
| --- | --- | --- | --- | --- | --- |
| G62R | rs761501352 | T65R | + | + | ATP binding |
| R93Q | rs781779034 | R96Q | +/- | - | ATP binding |
| R93G | rs28756978 | R96G | ND | - | ATP binding |
| P183L | rs759067670 | P199L | +/- | - | ATP binding |
| N291K | rs767413852 | L313K | + | + | Conserved amino acid |
| F390I | rs61752721 | R407I | +/- | + | undefined |
| R1230C | rs746431837 | R530C | - | - | Endonuclease |
| R1230H | rs781739661 | R530H | - | - | Endonuclease |
| R1232C | rs550698696 | R532C | +/- | + | Endonuclease |
| A1394T | rs138006166 | A702T | +/- | - | Endonuclease |
| G1396W | rs368876363 | G704W | +/- | +/- | Endonuclease |

**Legend** – “Variant” refers to amino acid change conferred by the indicated SNP. “Yeast equiv” is the orthologous amino acid in *Saccharomyces cerevisiae*. MMR, mismatch repair phenotype. CO, crossing over phenotype. For these phenotypes, “+” is WT level, “-“ is null, and “+/-“ is intermediate. ND, no data. See Table S3 for the complete dataset.
