## Supplementary Table 3 for "*MLH1/3* variants causing aneuploidy, pregnancy loss, and premature reproductive aging"

**Table S3a. Analysis of yeast equivalents to human *mlh3* SNPs**

---*lys2-A_14_* reversion rate---	**Phenotype**

***mlh3* allele	(x 10^-6^)	95% CI (x 10^-6^)	n	relative to WT	% tetratype (n)	relative to WT	MMR	CO**

*MLH3*	3.70	1.27-6.80	14	1	39.9 (1052)	1	+	+

*MLH3::KANMX^#^*	39.8 (1004)	1.0	+

*mlh3Δ*	23.1	15.1-29.4	16	6.24	18.4 (706)	0.49	-	-

*T65R*	4.60	3.51-10.8	11	1.24	36.4 (272)*	0.97	+	+

*R96Q*	11.9	8.75-16.7	16	3.21	22.8 (302)**	0.61	+/-	-

*R96G*	not tested	21.8 (1122)**	0.58	-

*P199L*	12.9	9.77-17.8	16	3.49	23.1 (312)**	0.62	+/-	-

*L313K*	6.34	4.24-9.29	11	1.71	31.2 (266)*	0.83	+	+

*R407I*	12.7	5.20-17.3	16	3.43	38.9 (841)*	1.04	+/-	+

*R530C*	18.1	11.2-29.1	11	4.89	21.3 (258)**	0.57	-	-

*R530H*	17.7	13.1-19.2	11	4.78	19.9 (246)**	0.53	-	-

*R532C*	10.1	7.37-13.8	16	2.73	36.5 (537)*	0.97	+/-	+

*A702T*	9.06	6.63-10.9	16	2.45	24.6 (666)**	0.66	+/-	-

*G704W*	8.64	6.82-14.6	11	2.34	30.4 (642)***	0.81	+/-	+/-

**Legend -** *lys2-A_14_* reversion (n= number of independent assays) and meiotic crossover analyses. For % tetratype (two spores containing parental, and two containing nonparental information, indicative of a crossover event), n represents the sum of parental ditype and tetratype events. For phenotypes: +, wild-type; -, null; +/-, intermediate. *p<0.0001 compared to null, p>0.002, compared to wild-type; **p>0.002 compared to null, p<0.0001 compared to WT; ***p<0.0001 compared to null, p<0.0001 compared to WT. A Bonferroni correction was applied for the Pearson Chi-Squared contingency test of the meiotic crossover data. 26 comparisons were made (13 to *MLH3::KANMX*, 13 to *mlh3Δ*), for a p<0.002 (0.05/26) cut off for statistical significance. ^#^ No difference in the reversion rate at the *lys2A_14_* locus was observed between *MLH3* and *MLH3::KANMX* strains (Nishant et al., *Genetics* 179: 747, 2008).

**Table S3b Statistics for meiotic crossover assay in Table S3a**

|  | ***mlh3* allele** | | **PD** | | **TT** | | **PD+TT** | | **%TT** | | **Pvalue to *MLH3::KANMX*** | | **Pvalue to null** | | **Phenotype** |
| --- | --- | --- | --- | --- | --- | --- | --- | --- | --- | --- | --- | --- | --- | --- | --- |
|  | *MLH3* | | 635 | | 417 | | 1052 | | 39.9 | | 0.92 | | <0.0001 | | + |
|  | ***MLH3::KANMX*** | | 604 | | 400 | | 1004 | | 39.8 | | **1** | | <0.0001 | | + |
|  | ***mlh3null*** | | 576 | | 130 | | 706 | | 18.4 | | <0.0001 | | **1** | | - |
|  | T65R | | 173 | | 99 | | 272 | | 36.4 | | 0.30 | | <0.0001 | | + |
|  | R96G | | 877 | | 245 | | 1122 | | 21.8 | | <0.0001 | | 0.078 | | - |
|  | R96Q | | 233 | | 69 | | 302 | | 22.8 | | <0.0001 | | 0.11 | | - |
|  | P199L | | 240 | | 72 | | 312 | | 23.1 | | <0.0001 | | 0.085 | | - |
|  | L313K | | 183 | | 83 | | 266 | | 31.2 | | 0.0099 | | <0.0001 | | + |
|  | R407I | | 514 | | 327 | | 841 | | 38.9 | | 0.67 | | <0.0001 | | + |
|  | R530C | | 203 | | 55 | | 258 | | 21.3 | | <0.0001 | | 0.31 | | - |
|  | R530H | | 197 | | 49 | | 246 | | 19.9 | | <0.0001 | | 0.60 | | - |
|  | R532C | | 341 | | 196 | | 537 | | 36.5 | | 0.20 | | <0.0001 | | + |
|  | A702T | | 502 | | 164 | | 666 | | 24.6 | | <0.0001 | | 0.005 | | - |
|  | G704W | | 447 | | 195 | | 642 | | 30.4 | | <0.0001 | | <0.0001 | | +/- |

P-value: Pearson Chi-Squared contingency test, Vassar Stats, with a Bonferroni correction for 26 comparisons, for a p<0.002 cut off for statistical significance.
