## Supplementary Table 4 for "*MLH1/3* variants causing aneuploidy, pregnancy loss, and premature reproductive aging"

**Table S4. Yeast strains used in this study.**

| **Strain** | **Genotype** | | **Plasmid** |
| --- | --- | --- | --- |
| EAY3252 | *MATα, ho::hisG, ura3, leu2::hisG, trp1::hisG, ADE2, HIS4, CEN8Tomato::LEU2, MLH3, lys2::insE-A_14_* | |  |
| EAY4480 | *MATα, ho::hisG, ura3, leu2::hisG, trp1::hisG, ADE2, HIS4, CEN8Tomato::LEU2, MLH3::KANMX, lys2::insE-A_14_* | |  |
| EAY3255 | *MATα, ho::hisG, ura3, leu2::hisG, trp1::hisG, ADE2, his4xB, CEN8Tomato::LEU2, mlh3Δ::NATMX, lys2::insE-A_14_* | |  |
| EAY3486 | *MATa, ho::LYS2; lys2; ura3; leu2::hisG; trp1::hisG; THR1::m-Cerulean-TRP1; mlh3Δ::NATMX* | |  |
| EAY3944-3945 | Same as EAY3255, but *mlh3-R530C::KANMX* | | pEAI409 |
| EAY3946-3947 | Same as EAY3255, but *mlh3-R530H::KANMX* | | pEAI410 |
| EAY3948-3949 | Same as EAY3255, but *mlh3-L313K::KANMX* | | pEAI411 |
| EAY3950-3951 | Same as EAY3255, but *mlh3-T65R::KANMX* | | pEAI412 |
| EAY3953, 3982, 3983 | Same as EAY3255, but *mlh3-R96Q::KANMX* | | pEAI413 |
| EAY4133, 4134 | Same as EAY3255, but *mlh3-R96G::KANMX* | | pEAI441 |
| EAY3954-3955 | Same as EAY3255, but *mlh3-R532C::KANMX* | | pEAI414 |
| EAY3956-3957 | Same as EAY3255, but *mlh3-G704W::KANMX* | | pEAI415 |
| EAY3958-3959 | Same as EAY3255, but *mlh3-R407I::KANMX* | | pEAI416 |
| EAY3960-3961 | Same as EAY3255, but *mlh3-A702T::KANMX* |  | pEAI417 |
| EAY3979-3981 | Same as EAY3255, but *mlh3-P199L::KANMX* |  | pEAI418 |

All strains are from the SK1 background. “Plasmid” refers to the *mlh3::KANMX* vector used to integrate the indicated *mlh3* allele.
